# Diverse transcriptomic response of cellular system following low-dose exposure of mesoporous nanoparticles

**DOI:** 10.1101/2022.09.16.508239

**Authors:** Deepti Mittal, Syed Azmal Ali

## Abstract

Mesoporous nanoparticles (NPs) are an interesting drug delivery system that has generated considerable attention in the biomedical sector. Despite recent attempts to conduct safety assessments using traditional methods based on phenotypic data, our understanding of the underlying molecular processes produced by mesoporous NPs is still in its infancy. In the present study, RNA sequencing was used to assess the biological perturbations and the pathways induced in response to early exposure of two different mesoporous NPs; mesoporous silica NPs (MSN) and mesoporous carbon NPs (MCN) in human liver hepatocellular carcinoma cells. In order to better understand the risks associated with NPs, it is required to consider the initial low-dose exposure effects that mimic the real exposure scenario. No overt toxicity was detected in the MTT assay when performed at 6 hours at low concentrations (MCN 25 g/ml and MSN 15 g/ml) of NPs; thus, we have selected this dose for RNA sequencing analysis. Our transcriptomics analysis showed significant differences in the expression of many genes after exposure to both NPs. Surprisingly, both NPs frequently deregulated 52.9 percent of upregulated and 42 percent of downregulated genes. Gene ontology categories, in particular, revealed comparable perturbations of biological reactions in the cellular system. HepG2 cells reacted to mesoporous NPs by allowing alterations in genes involved in cytoskeleton reorganisation (ATAT1, DMTN, PTK2 and PFN2). Exposure to mesoporous NPs increased transcripts expressing ubiquitin ligase (RNF187, ARIH2, VHL, and RAB40C), transferase (FBXO3 and WDSUB1), conjugating (UBE2J2), and also proteasomal subunits (PSMD2, PSMD13) enzymes, indicating that protein turnover rates are altered in response to environmental damage. In addition, DNA damage and DNA damage checkpoint genes were upregulated, indicating that NPs induced stress in the cells. These finding showed low dosage acute exposure have comparable responses between mesoporous NPs. These results may add further knowledge in conceptualization of Safer-by-Design strategy of NPs in biomedical field.

## 1. Introduction

The nanoparticles (NPs) are novel materials compared to their bulk counterparts, owing to their unique physical and chemical properties. In the present study, we have considered two important mesoporous NMs: mesoporous silica nanoparticles (MSN) and mesoporous carbon nanoparticles (MCN). According to the International Union of Pure and Applied Chemistry (IUPAC), mesoporpous materials are defined as materials with pore diameters ranging from 2 to 50 nm. The highly ordered, porous, tunable and textural properties make them a suitable candidate for targeted drug delivery applications (Mittal and Ali, 2022; Manzano and Regi, 2010), tissue regeneration (Chen et al., 2019) and biosensors (Ali et al., 2021; Cho et al., 2020). In addition, considerable attention has been paid towards the environmental application of these highly porous and thermally stable materials as a removal agent for environmental pollutants (Cashin et al., 2017; Zhang et al., 2019). The increased utilisation of these NPs leads to an unrecognised environmental exposure that can promote consequences for general public health and raise environmental concerns. Therefore, these NPs must be thoroughly studied in order to gain sufficient knowledge about their potential impact. However, we are still unclear how these NPs interact with the cellular milieu and cause the potential impact on human health that must be assessed to produce safer materials (Vishwakarma et al., 2010). There are different schools of thought regarding the safety of mesoporous NPs, due to which the safety issue of these NPs is still in its infancy. Few mechanistic studies have been carried out to find the safety profile of mesoporous nanostructures. MSN induced reactive oxygen species (ROS) and apoptosis followed by 24 h exposure in intestinal epithelial cells (Deng et al., 2021). Heikkila and coworkers reported ROS production at doses < 1 mg/ml in keratinocyte cell line and not at the low doses (Heikkilä et al., 2010). MSN also caused significant upregulation of the inflammatory genes through expression of NF-κB and AP-1 (Chou et al., 2017). MCN also induced toxicity by impairing cellular acivity of lung epithelial cells and triggered macrophage activation (Chen et al., 2017). Our lab has recently demonstrated the effect of MSN on proteome profile of CHO-K1 cell line and found many pathways related to Rho-Rac signalling that ultimately affects cytoskeleton of the cell (Yadav et al., 2021). Other than this we have also conducted transcriptomics analysis to study the effect of ZnO NPs and SWCNT in liver cell line that particularly provided clues about the general stress response induced by the NPs. Both the NPs induced cell cycle machinery, DNA damage and repair, and also affected ubiquitination pathway during the initial (acute) interaction with the cell (Mittal et al., 2022). Therefore, it is important to further characterize these NPs at a molecular level for their safer utilization.

The majority of studies, based on the safety analysis of NPs, relies exclusively on a single endpoint assay that includes cell viability, DNA damage response (Duan et al., 2013), apoptosis and necrosis (Mohammadinejad et al., 2019), and inflammation (Khanna et al., 2015). However, these single endpoints are not necessarily depicting the actual nano-bio interactions at the molecular level, as there is a possibility that molecular changes might have occurred before phenotypic changes. In this regard, omics techniques play an essential role in predicting the actual changes at the deeper molecular level, which could serve as a new target or a biomarker for particular NPs. Additionally, the hazard assessment should be foremost as the initial strategy in the conceptualization of Safer-by-Design approach that may enable us to select the most promising drug delivery candidate.

Prior to now, the majority of safety evaluation studies using NGS relied on high dosage exposure circumstances that did not represent a realistic situation. Understanding the toxicological consequences of each kind of NP is essential for estimating their short- and long-term dangers to people and ecosystems. Determining the low-dose and non-cytotoxic effects of MSN and MCN NPs in order to get a deeper understanding of their interaction with cells was the objective of our investigation.

## 2. Material and Methods

### 2.1. Nanoparticles suspension preparation

The NPs synthesis and characterisation was previously described by Rawat N. et al. 2016, and Yadav et al. 2021. The NPs were diluted in phosphate buffer saline (PBS). The PBS was used to prevent any unwanted interaction of NPs that could be possible with cell culture media components. The stock solution of NPs was sonicated by ten cycle pulse (pulse ON 15 s and OFF 15 s) on ice to avoid excessive heating and agglomeration.

### 2.2. Cell culture and exposure of nanoparticles

HepG2 cells were procured from National Centre for Cell Sciences (NCCS), Pune, India, and grown in DMEM 1X medium supplemented with 10% FBS, 1% streptomycin and penicillin solution, in 25 cm^2^ flask, at 37°C in 5% CO_2_ humified atmosphere. The cells were used between 15-20 passages for each experiment. HepG2 cells were plated at a density of 3×10^5^ cells/ml in a 96 well plate and allowed to attach for 24 h before the treatment of NPs. The NPs were suspended in PBS to make the 5 mg/ml stock solution and diluted to desired concentrations. The stock solution of NPs was sonicated on an ice bath for 10 min before treating the cells using a Branson ultrasonicator with a 13 mm probe diameter.

### 2.3. MTT assay

The cell viability was assessed by utilising the MTT assay according to Mosmann (1983) method with some modifications (Mosmann, 1983). First, the HepG2 cells were exposed to the NPs at the indicated concentrations for the different incubation periods. The medium was then removed and replaced with 10 μl of MTT [3-(4, 5-dimethylthiazoyl-2-yl)-2,5-diphenyltetrazolium bromide] dye (5 mg/ml) prepared in PBS and incubated for 4 h at 37°C. After incubation, the resulting formazan crystals were dissolved by adding 100 μl of dimethylsulphoxide, and the absorbance was measured at 570 nm in a multiplate reader (BioTek Instruments, Winooski, Vermont, USA). Three independent experiments were performed.

### 2.4. WST-8 assay

The final product of WST-8 is more soluble than MTT formazan in cell culture media and has higher sensitivity than other tetrazolium salts (Tominaga et al., 1999; Chamchoy et al., 2019), thus selected for further viability assessment. The WST-8 assay kit was used for NPs mediated cell viability analysis, following manufacture protocol. The HepG2 cells were seeded at a density of 2×10^5^ cells per ml in 96 well plate, allowed to adhere for 24 h and exposed to different concentrations of MCN and MSN NPs. After 24 h, 10 μl of WST-8 were added to each NPs treated well and incubated at 37° C for 2 h. The absorbance was then measured at 450 nm using a multi-plate reader (BioTek Instruments, Winooski, Vermont, USA).

### 2.5. Cell exposure for RNA sequencing experiment

HepG2 cells were allowed to grow for 24 h, at a density of 10^6^ cells/ml, in a 25 cm^2^ cell culture flask. Then, the medium was replaced with the fresh cell culture medium containing MCN and MSN. Finally, cells with only a fresh cell culture medium were used as a control sample. Low cytotoxic concentrations were chosen (MCN: 25 μg/ml, MSN: 15 μg/ml) for further transcriptomic analysis at which more than 80% viability was achieved (according to cell viability assay).

### 2.6. RNA purification, library construction and RNA sequencing

Total RNA was extracted from nanomaterial treated and untreated (control) HepG2 cells using the RNeasy mini kit (Qiagen GMBH, Hilden, Germany) following the manufacturer’s recommendations. The integrity of total RNA was assessed using the Agilent 2100 bioanalyzer (Agilent Technologies, Santa Clara, CA, USA). The NEBNext® Ultra™ Directional RNA Library Prep Kit (New England Biolabs, MA, USA) was used to prepare the RNA-seq library of an individual sample. The average RIN was >9.0 (Supplementary figure 1), and the 1 μg of pooled total RNA was taken for mRNA isolation, fragmentation and priming. Fragmented and primed mRNA was further subjected to the first-strand synthesis in Actinomycin D (Gibco, Life Technologies, CA, USA), followed by second-strand synthesis. The double-stranded cDNA was purified using HighPrep magnetic beads (Magbio Genomics Inc, USA). Purified cDNA was end-repaired, adenylated and ligated to Illumina multiplex barcode adapters. Adapter ligated cDNA was purified using HighPrep beads and was subjected to 12 cycles of Indexing-PCR (37 °C for 15 mins followed by denaturation at 98°C for 30 sec, cycling (98°C for 10sec, 65°C for 75sec) and 65°C for 5mins) to enrich the adapter-ligated fragments. The final PCR product (sequencing library) was purified with HighPrep beads, followed by a library-quality control check. The Illumina-compatible sequencing library was quantified by Qubit fluorometer (Thermo Fisher Scientific, MA, USA), and its fragment size distribution was analysed on Agilent 2200 Tapestation (Supplementary figure 2, A-I). Samples were then sequenced using the Illumina Genome Analyzer (HiSeq 2500v4 High Output).

### 2.7. Data analysis

FastQC (version 0.11.2) was used for quality control to ensure that the quality value was above Q30. The Human (*Homo sapiens*) genome FASTA file (ftp://ftp.ensembl.org/pub/release-100/fasta/homo_sapiens/dna/) and gene annotation GTF file (Human (GRCh38.p13) assembly; ftp://ftp.ensembl.org/pub/release-100/gtf/homo_sapiens/) were obtained from Ensembl. Although RNA-seq is a popular research tool, there is no gold standard for analysing the data. Therefore, we chose up-to-date open source tools for mapping, retrieving read counts, and differential analysis among the available tools. We used HISAT2 (version 2.0.5) (Kim et al., 2015) to generate indexes and map reads to the human genome. For assembly, we chose SAMtools (version 1.2) and the “union” mode of HTSeq (version 0.6.1) (Anders et al., 2015), as the gene-level read counts could provide more flexibility in the differential expression analysis. Both HISAT2 and HTSeq analyses were conducted in the Linux operating system (version 2.6.32). Differential expression and statistical analysis were performed using DESeq2 (release 3.3) in R (version 3.2.4). DESeq2 (Love et al., 2014) was chosen as it is a popular parametric tool that provides a descriptive and continually updated user manual (https://bioconductor.org/packages/release/bioc/vignettes/DESeq2/inst/doc/DESeq2.html).

DESeq2 internally corrects library size, so it is crucial to provide un-normalised raw read counts as input. We used variance stabilising transformation to account for differences in sequencing depth. P-values were adjusted for multiple testing using the Benjamini-Hochberg procedure (Benjamini and Hocheberg, 1995). A false discovery rate adjusted p-value (i.e., q-value) < 0.05 was set to select DE genes.

### 2.8. Bioinformatics analysis

The list of differential expression of genes was utilised to observe transcriptional snapshots through bioinformatics analysis. The GO and pathway were initially identified through DAVID analyses Overrepresentation Test (released on July 15, 2016) in PANTHER version 11.1 (http://www.pantherdb.org/, released on October 24, 2016). This program supports the canine genome. PANTHER used the binomial test and Bonferroni correction for multiple testing, while DAVID used the right-tailed Fisher Exact test and displayed z-scores. Only pathways and GO terms with fold enrichment > 0.2 and p-value > 0.05 were listed in PANTHER. The pathway enrichment analysis of up and downregulated genes was performed using the R package, cluster profile package (Yu et al., 2012) with a p-value cutoff ≥of 0.05. All the graphical analyses were performed in the R environment using respective tools. Volcano plots and Principal Component Analysis (PCA) were generated using the ggPlot2 package. Protein-protein interaction networks were created using Cytoscape

### 2.9. Functional Annotation Analysis

We performed (Database for Annotation, Visualization and Integrated Discovery) DAVID analysis (Dennis et al., 2003) to obtain a comprehensive set of functional annotation of mesoporous nanostructures in a liver-based HepG2 cell line. Initially, we selected transcripts more than and less than 3 fold change differentially regulated and submitted them to DAVID web-based bioinformatics tool to gather information. Then, we performed the DAVID as it functionally annotates clusters for the identified terms. The newly updated enrichment analytic algorithm measure the relationships among annotation terms based on the degrees of their co-association genes to group similar, redundant, and heterogeneous annotation contents from the same or different resources into annotation groups. We calculated the fold enrichment by keeping the minimum number of 2 genes for the corresponding term and maximum EASE threshold/p-value of 0.1 using the multiple post-test including Bonferroni, Benjamini, and Fisher Exact test to increase confidence the annotation. We also calculated the corrected FDR values for all the tests mentioned above to reduce the burden of associating similar redundant terms and make the biological interpretation more focused on a group level. All the calculations for functional annotation clustering were performed in the background of the Homo sapiens genome database.

### 2.10 Statistical analysis

All the data were analysed using GraphPad Prism 5.01 (GraphPad Software, San Diego, California, USA, https://www.graphpad.com/). The significance of the difference between mean values was calculated by one-way analysis of variance (ANOVA) with Tukey’s post hoc test. The data were represented as mean ± SD, and the value of p < 0.05 was considered statistically significant.

## 3. Results and Discussion

### 3.1 Cytotoxicity measurement and dose selection for RNA sequencing

NPs effects have been shown to depend upon the dose delivered to cells (Efeoglu et al., 2016). Therefore, it becomes necessary to select the appropriate NPs dose to understand the actual scenario in environmental settings. To determine the optimal concentration of MCN and MSN NPs for transcriptomics analysis, cell viability was determined 3, 6, and 12 hours after treatment with various dosages of MCN and MSN NPs (Figure 1).

**Figure 1:**
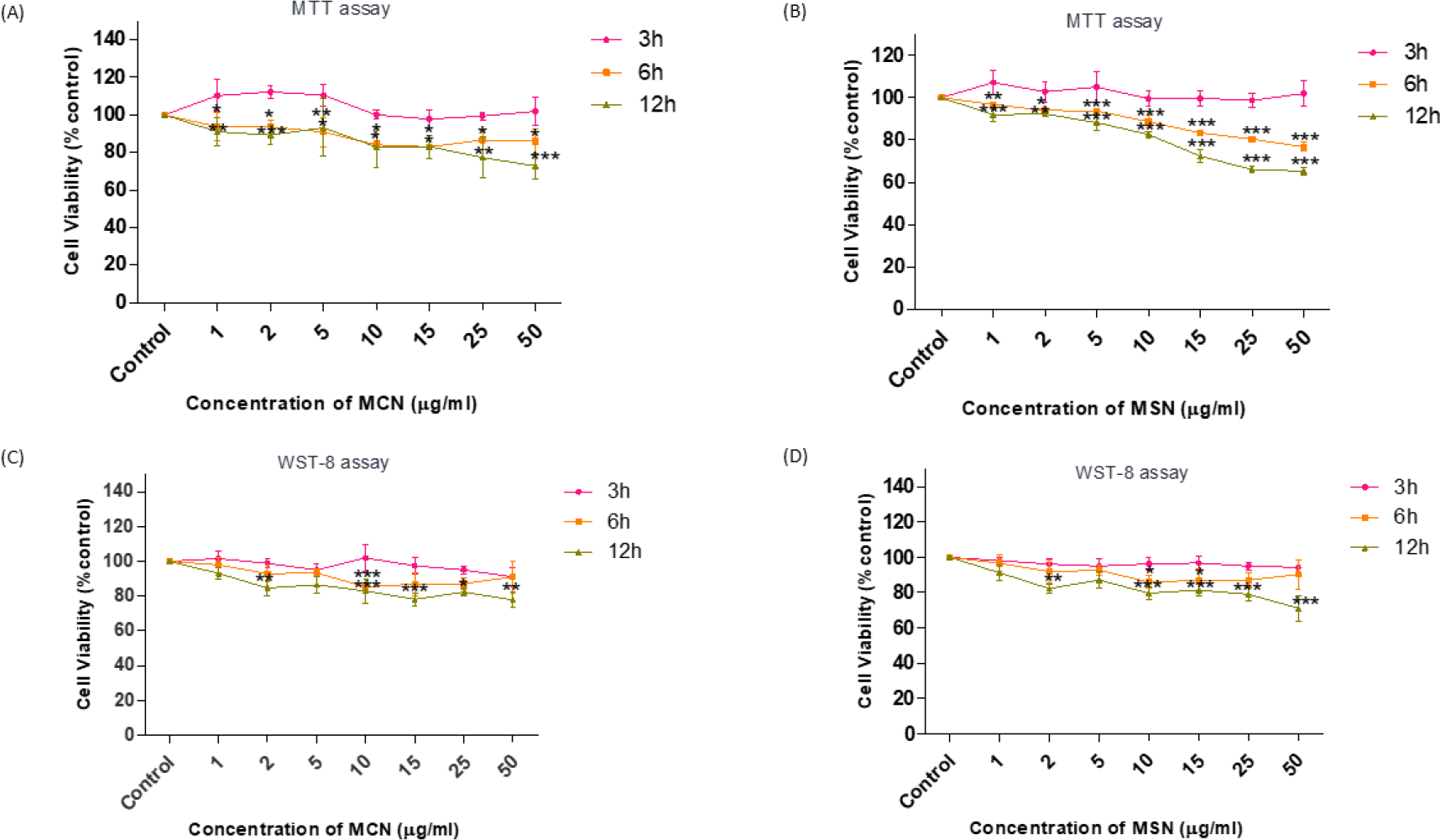
Preliminary cell compatibility analysis of MCN and MSN NPs: Cell cytotoxicity analysis in HepG2 cells was conducted by MTT (A, B) and WST-8 (C,D) assay after the exposure of MCN and MSN NPs for 3, 6, and 12 h treated with different concentrations (1, 2, 5, 10, 15, 25, 50 μg/ml). All the assay values were normalized with non-treated control, the viability of which is represented as 100%. Results are shown as mean values ± SD obtained from three independent experiments (n=3). Statistical analysis was performed using two-way ANOVA followed by Bonferroni post-test.

During the initial incubation period of 3 h, both the NPs are non-cytotoxic on the HepG2 cells even at the higher dose of 50 μg/mL. Surprisingly, increased viability was observed at 3 h, suggesting the increased metabolic activity of cells with the NPs exposure as the MTT assay is used to measure the metabolic redox activity; therefore, the sudden load prepared the cells to deal with the NPs.

On the other hand, compared to the MCN the MSN NPs are diminishing the viability of cells at higher concentrations depicting in both MTT and WST-8 assay. The MTT assay results showed decreased cell viability to 28% at 15 μg/mL at the incubation period of 12 h, unlike the MCN that decreased the 17 % viability at a similar concentration at 12 h (Figure 1A and B). This is in agreement with a recent publication from our lab that suggested 15 μg/mL as the Low Observed Adverse Effect Level (LOAEL) at 24 h exposure period compared to other NPs (ZnO NPs and MWCNT) (Yadav et al., 2021). Here we compare the effects of two mesoporous NPs with different compositions; Carbon in MCN and Silica in MSN. Very few studies considered MCN for cytotoxicity analysis (except few studies considering MCN for drug delivery applications) compared to MSN. Thus, our study provides initial and valuable information about their biocompatibility.

Similar results were found in the case of WST-8 assay where >90 % of the viability was shown by cells treated with MCN and MSN at 3 h at all the considered doses (Figure 1C and D). However, significant (p<0.01) viability loss was seen at 50 μg/mL of dose of MCN and MSN also at 12 h incubation period. At the 6 h exposure period, the cell viability was maintained at >80% at all the considered concentrations. Therefore, this period was selected further to explore the molecular changes. The dose of 15 μg/mL was considered for MSN (according to the previously published paper from our lab, Yadav et al., 2021) and 25 μg/mL for MCN NPs. At this dose, >85% cell viability was maintained.

#### 3.2 Transcriptional tour of HepG2 cells exposed to two different mesoporous structures

The main advantage of employing high-throughput techniques such as RNA sequencing is to extract the global gene expression responses without a *priori*, despite collecting information in a hypothesis-driven manner. In the present study, we employed high-throughput RNA sequencing to analyse deeply the molecular changes associated with the NMs insult (Figure 2).

**Figure 2:**
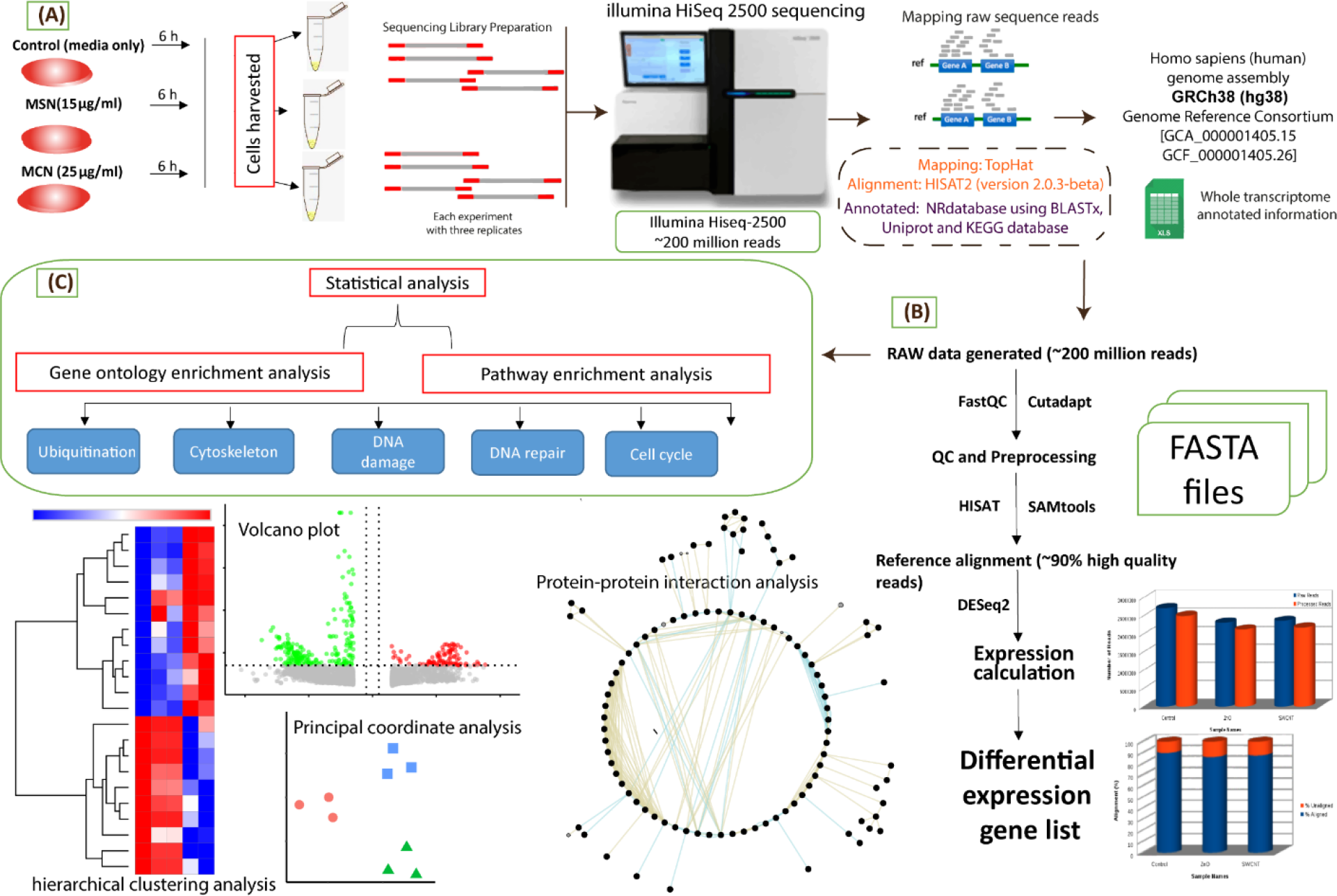
(A) Systematic transcriptomics workflow representing exposure of HepG2 cells with MCN and MSN NPs followed by the RNA extraction and subsequent sequencing of samples through Illumina Hiseq 2500 platform. (B) Raw sequencing data was trimmed, quality checked and further aligned to the reference genome of the human (GRCh38). DESeq2 was utilized for expression calculation to get the list of differentially expressed genes (DEG). (C) The DEG list was utilized to carry out the enrichment analysis that significantly provided similarly enriched terms for both the NPs.

The genome-wide gene expression analysis was performed on Illumina Hiseq 2500 (150*2 chemistry), generating about ~87 million high-quality adapter free reads in all the samples. The generated raw data has been checked for quality using FastQC and pre-processed using Cutadapt, which includes removing adapter sequences and the low-quality bases (<q30) (Figure 2A & B). On average, 29.17 million reads were retained for further downstream analysis. An average of 94.99% of the high-quality reads were aligned to the Homo sapiens reference genome GRCh38 using HISAT2, a fast and sensitive alignment program for mapping next-generation sequencing reads (Supplementary figure 3). The total number of raw reads was distributed among the three analysed samples: control 29809925, MCN 29327469, and MSN 33954023. After the normalisation, the transcripts were identified and quantified using Cufflinks based on the obtained aligned reads. Cufflinks were used to calculate the differentially expressed transcripts and categorise them into up, down and neutrally regulated genes based on the log2 fold change values. In the control sample, 87501 transcripts were expressed, and in the case of MCN and MSN, the numbers were 88294 and 89527, respectively. The expression was measured in FPKM (Fragments Per Kilobase of transcript per Million mapped reads) units for every transcript. The number of reads corresponding to the particular gene is normalised to the length of the gene and the total number of mapped reads. The PCA shows the distinct pattern expression profile in MSN and MCN treated cells (Figure 3).

**Figure 3:**
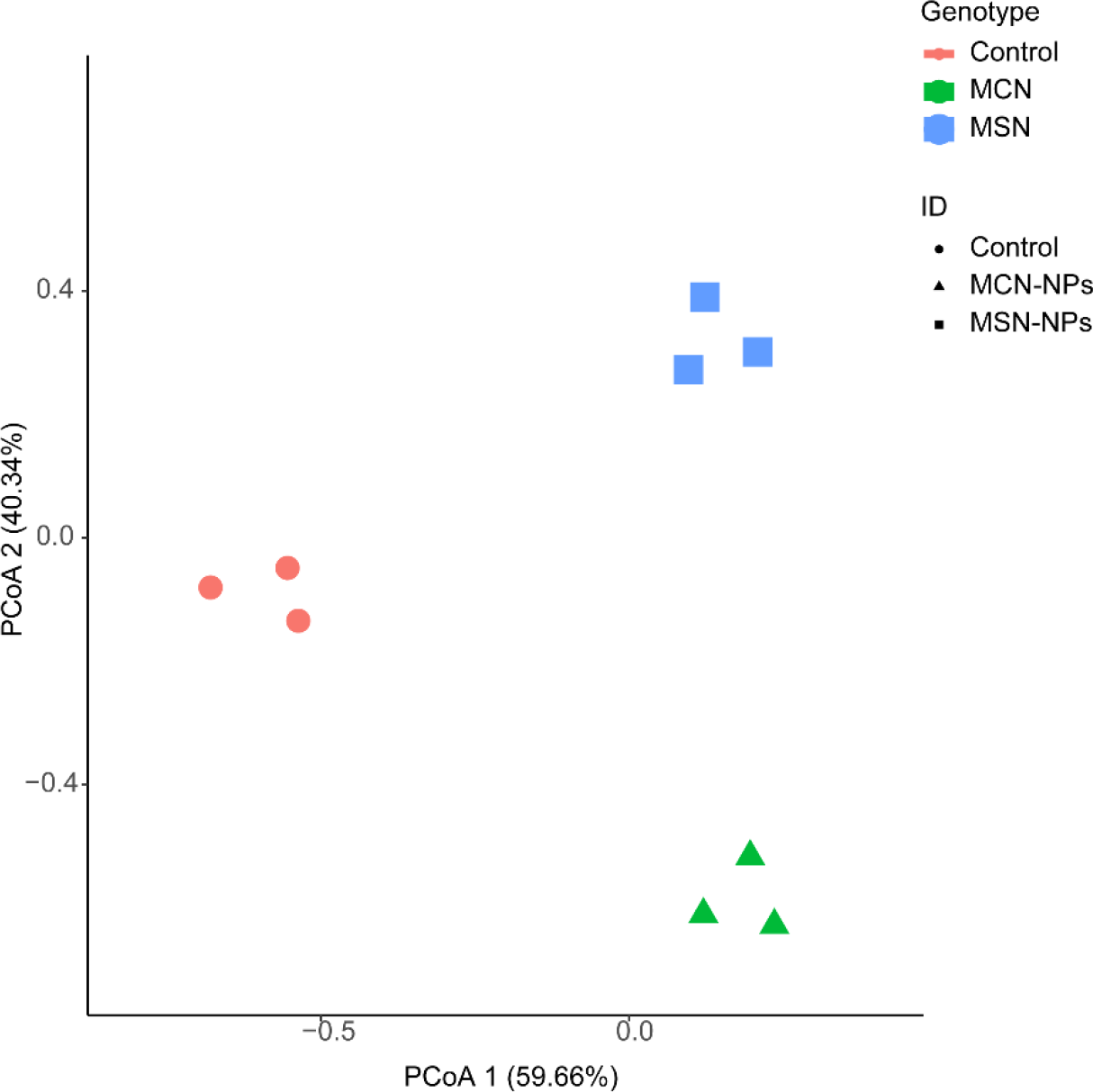
The PCA plot showing the disctribution of transcriptomics data

Next, sample wise comparison was performed to identify differentially regulated transcripts between two mesoporous structures MSN and MCN. Transcripts with expression values less than 0.66 are considered down-regulated, while higher than 1 are considered upregulated, and those with an expression value in the range of 0.66 to 1 are considered neutrally regulated. We have emphasised the unbiased analysis of high-throughput studies to gain deeper insights even with the lower expressed transcripts that might give the idea of essential biomarkers of exposure. The number of differential expression of genes (non-redundant) for MCN was 10090 (5859 upregulated and 4231 downregulated) and 10179 for MSN (6188 upregulated and 3991 downregulated) (Supplementary figure 4). Surprisingly, 52.9% (4166) upregulated and 42% (2433) downregulated genes were shared between MCN and MSN, providing clues for the similar molecular mechanisms followed by mesoporous NPs with the differing core. The volcano plot showed the differentially abundant transcript between MCN and MSN (Figure 4). This is the first study to date comparing the cellular effects of MCN and MSN at the molecular level at non-cytotoxic doses. The primary reason for comparing these NPs relies on their potential role as a drug delivery vehicle (Zhao et al., 2017; Manzano and Regi, 2020); thus, the study of their interaction with cells will provide important information useful for direct Safe-by-Design strategy.

**Figure 4.**
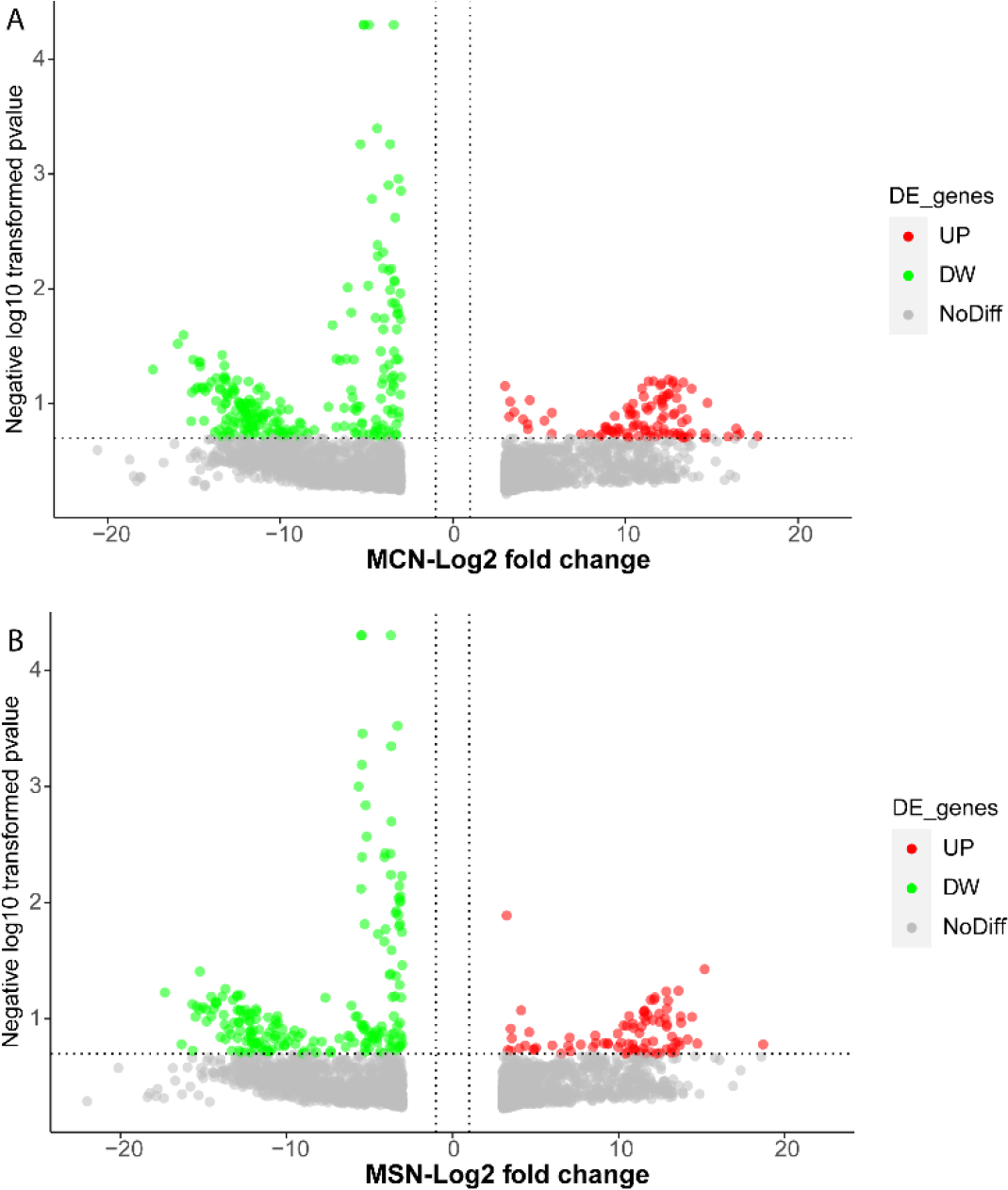
Volcano plots comparing the differentially expressed transcripts in comparision to control cells.

#### 3.3 Ontology-driven approach to find significant terms for NPs exposure

By using various databases, we performed functional characterization of differentially expressed genes (DEGs) among the GO categories, biological process (BP), molecular function (MF) and cellular component (CC), to observe the critical response. We selected significant (p≤0.05) ontological terms with multiple hypothesis 2 post-test corrected p values (p≤0.05) Bonferroni and (p≤0.05) Benjamini corrected. The enriched biological function for upregulated genes against MCN comprised of cell-cell adhesion (GO:0098609), protein ubiquitination (GO:0016567), DNA damage response (GO:0006977), DNA repair (GO:0006281), phosphorylation (GO:0006468), mRNA splicing via spliceosome (GO:0000398), translation initiation (GO:0006413) and MAPK cascade (GO:0000165). Similarly, downregulated genes encompassed BP categories, including xenobiotic metabolic process (GO:0006805), DNA replication (GO:0006260), protein phosphorylation (GO:0006468), negative regulation of protein ubiquitination (GO:0031397) and protein ubiquitination (GO:0016567). The DEGs regulated by the MSN significantly enriched for protein polyubiquitination (GO:0000209), DNA repair (GO: 0006281), proteasome-mediated ubiquitin-dependent protein catabolic process (GO:0043161), cell-cell adhesion (GO:0098609) and regulation of apoptotic process (GO:0042981).

The significant annotated genes are found at different locations inside the cells and maximally enriched in the cytoplasm (428), followed by the nucleus (386) and cytosol (310). The expressed genes enriched for protein binding, protein kinase, ATP binding, poly(A) RNA binding, chromatin and zinc ion binding execute the following molecular activities. The GO is summarized in full in Table S2, containing all of the ontologies for MCN and MSN. We discovered some intriguing findings after doing a thorough analysis of the data. The most significant top ontologies were common for MCN and MSN (Supplementary table 3), which directs our hypothesis towards a similar response against non-cytotoxic doses at acute exposure. We have selected and focused on the highly significant ontology commonly shared by MCN and MSN.

#### 3.4 Cytoskeleton interaction with mesoporous nanoparticles and their modulation

Our RNA sequencing data also reflect the interaction with the cytoskeleton system and its potential uptake, giving clues for the efficient utilisation of mesoporous NMs as a drug delivery carrier. The Venn diagram represented 26% of upregulated and 20.4% downregulated genes with 2log FC≥3 commonly perturbed by both the NMs, thus providing evidence about a similar interaction pattern with cell cytoskeleton. We discovered that MCN had a 43.9% upregulation and a 41.2% downregulation of transcripts, while MSN had a 30.1% upregulation and 38.1% downregulation of genes (Figure 5A & B), which is critical for understanding the NPs unique effects.

**Figure 5:**
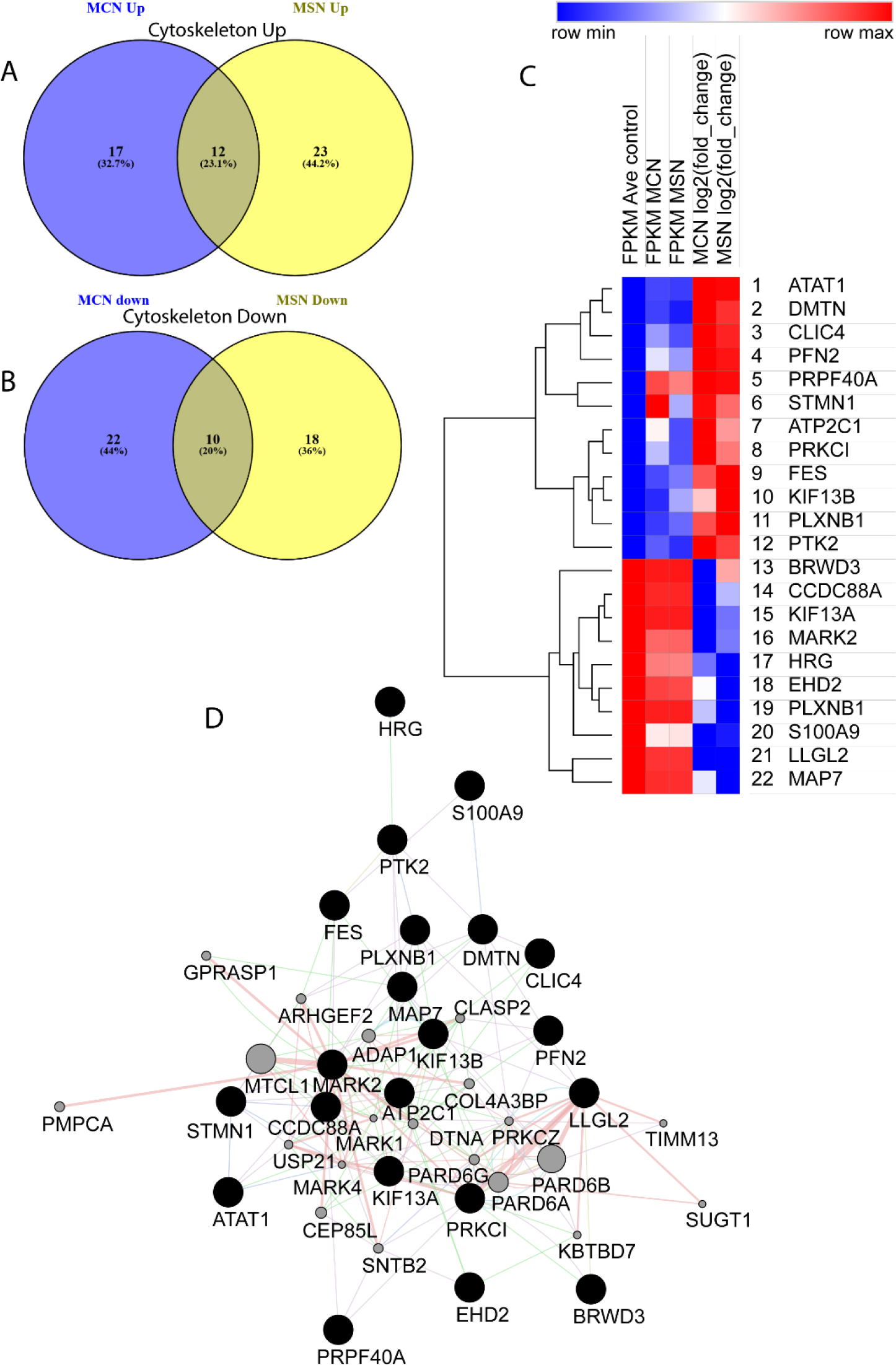
Cytoskeleton related gene alteration by mesoporous NPs: The Venn diagram represents unique and commonly shared deregulated genes of cytoskeletal machinery of the cell. (A) Represents upregulated genes regulated by MCN and MSN where 26% of the genes were common, 43.9% genes were regulated by MCN and 30.1% were regulated by MSN uniquely. MCN and MSN NPs commonly downregulated 20.6% genes whereas 41.2% genes by MCN and 38.1% genes by MSN NPs were uniquely downregulated. (C) Displaying heatmap of the genes detected in the overlap of the Venn diagrams regulated commonly by both the NPs. Columns are ordered by FPKM values of control, MCN and MSN NPs.

Under their interaction with the biological system, these NPs are capable of inducing cytoskeletal related changes as highlighted in the GO analysis that is commonly enriched with cell-cell adhesion (GO:0098609, MCN p-value 2.09E-05, MSN 5.99E-04) and regulation of cytoskeleton organisation (GO: 0051493, MCN p-value 0.412201154, MSN p-value 0.390015601) (Supplementary table 3).

Importantly, the cytoskeletal changes during NPs interaction signify their uptake and translocation through the cellular cytoplasm (Ispanixtlahuatl-Meráz et al., 2018) that further corroborates our transcriptomics data. Similarly in Molecular function the enriched category of cadherin binding involved in cell-cell adhesion (GO:0098641, MCN p-value 2.10E-05, MSN p-value 2.37E-04) with subsequent cellular component category terms including focal adhesion (GO:0005925, MCN p-value 3.04E-06, MSN p-value 0.00147841), cell-cell adherens junction (GO:0005913, MCN p-value 6.95E-06, MSN p-value 1.19E-04), actin cytoskeleton (GO:0015629, MCN p-value 0.003725131, MSN p-value 0.025017052), and microtubule cytoskeleton (GO:0015630, MCN p-value 0.004113805, MSN p-value 0.015340609) (Supplementary table 2) provides strong correlation for NPs induced cytoskeletal remodeling. Corresponding cytoskeletal genes; *ATAT1, DMTN, CLIC4, PFN2, PRPF40A, STMN1, ATP2C1, PRKCI, FES, KIF13B, PLXNB1, PTK2* are among the upregulated genes and *BRWD3, CCDC88A, KIF13A, MARK2, HRG, EHD2, PLXNB1, S100A9, LLGL2, MAP7* among downregulated genes (Figure 5C) by 3 log2 fold change.

The upregulation of ATAT1 (alpha-tubulin-N-acetyltransferase1) responsible for the acetylation of tubulin signifies the changing microtubule that is an essential part of the cytoskeleton and plays an important role during cell polarisation, cell migration, invasion (Janke and Montagnac, 2017). Similarly, the profilin (PFN2) gene with dual actin and microtubule regulatory activity (Costa and Sousa et al. 2019) was shown to be upregulated that suggested cytoskeletal alterations with mesoporous NMs exposure. A previous paper from our lab has also reported the increased migration of CHO-K1 cells with MSN exposure, which results from cytoskeleton organization (Yadav et al., 2021). Mesoporous NPs to change the cell biomechanics can be exploited for many therapeutic applications such as cancer. Additionally, observing the interaction of NPs with cytoskeleton prevents cell disruption during cellular internalisation, and cargo can be delivered with ease. We have also seen the upregulation of the important protein-tyrosine kinase gene: PTK2 (MCN FC: 8.3, MSN FC: 10.9) that plays an important role in reorganization of cell cytoskeleton (Fabry et al., 2011). The protein-protein interaction (PPI) network (Figure 5D) also shows highly dense network with lethal giant larvae gene; LLGL2 (MCN FC: −11.12, MSN FC: −11.02) which is essential for cell division and cell migration is highly downregulated. LGL is a cytoskeletal protein containing many sites for phosphorylation and also maintains the cell polarity and proliferation (Russ et al., 2012). Another important protein kinase MARK2 found to be downregulated by NPs (MCN FC: −7.7, MSN FC: −5.4) and highly dense in the PPI network takes part in regulating the microtubule dynamics and cell polarity also takes part in the cell cycle activation and DNA damage response (Hubaux et al., 2015).

#### 3.5 DNA damage, repair and cell cycle process initiated by the exposure of mesoporous structures (MSN & MCN)

We have performed the RNA-sequencing studies early point, i.e. 6 h, to look into the molecular responses that are just initiated by the exposure of NMs. With the acute responses, we can understand the primary mechanism of NMs insult and decipher the functional processes elicited to maintain the homeostasis in the cell. Additionally, the cells generate a sequential response and repair pathways upon DNA damage to maintain homeostasis inside the cell (Sancar et al., 2004). Genes identified to be differentially regulated with corresponding log2 (fold change)>3 were subjected to DAVID gene ontology (GO) analysis. As a result, we obtained the GO terms related to DNA damage and repair that gives us the idea of mesoporous NMs to cause cellular disturbances. In GO analysis, the biological process related to DNA damage and repair such as DNA damage response (G0:0006977; MCN p-value 4.53E-04, MSN p-value 0.007577298) and DNA repair (G0:0006281; MCN p-value 9.86E-04, MSN p-value 3.99E-05) were significantly enriched for both the NPs (Supplementary table 3). Both the NPs are exerting similar expression of DNA damage and repair related genes which is evident from the venn diagram (Figure 6A).

**Figure 6:**
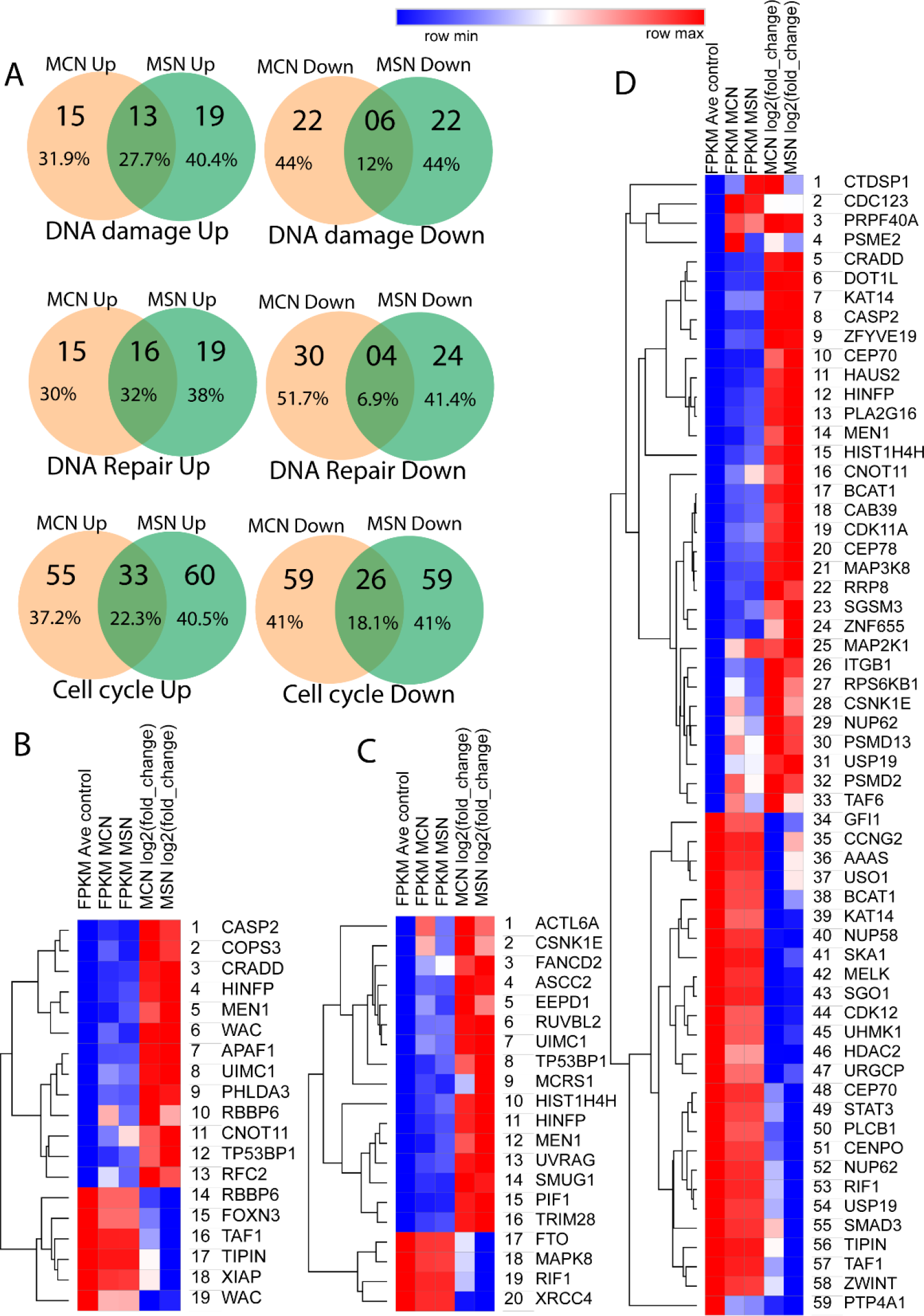
DNA damage, repair and cell cycle related gene alteration by mesoporous NPs: The Venn diagram represents unique and commonly shared regulated by both the NPs. (A) represents upregulated genes related to DNA damage regulated by MCN and MSN where 23.8% of the genes were common. DNA repair related genes commonly upregulated by MCN and MSN NPs are counted for 38.1% and downregulated are counted for 17.1%. Heatmap of the genes for DNA damage (B), DNA repair (C) and cell cycle (D) were detected in the overlap of the Venn diagrams regulated commonly by both the NPs. Columns are ordered by FPKM values of control, MCN and MSN NPs.

Notably, the genes such as Caspase-2; CASP2 (MCN FC: 9.8, MSN FC: 8.9) and CRADD (MCN FC: 8.4, MSN FC: 8.8) that takes part in the genotoxic stress response are consistently deregulated (Figure 6B). CASP2 is a vital initiator of caspase in apoptotic pathways and is known to be activated by the protein complex that includes death domain-containing protein, PIDD and CRADD (or RAIDD) (MCN FC: 8.4, MSN FC: 8.8) to form PIDDosome complex (Tinel and Tschopp, 2004). PIDD is expressed in response to the expression of p53, which was also seen to be upregulated (TP53; MCN FC: 7.9, MSN FC: 1.8) that further confirms the successful formation and activation of this complex. CASP2 is also responsible for releasing apoptogenic factors from the mitochondria when initiated by the external insult. It is also regarded as the direct activator of the apoptotic pathway (Guo et al., 2002). We have also seen the upregulation of APAF1 (MCN FC: 4.9, MSN FC: 4.10); Apoptotic protease activating factor-1, which also plays an essential role in regulating apoptosis and mediates the activation of other caspases (Shakeri et al., 2017), though not significantly enriched in our data. Thus, the activation of CASP2 and its effector molecules suggested that NPs has already started to kill the cells, or we can say that NPs has already initiated the pathways that might lead to apoptosis but not entirely kill the cells in this short duration of exposure (i.e. 6 h).

Interestingly, we have also enriched significantly the DNA repair pathways that might suggest the transient nature of the DNA damage or the damage imposed by the NPs primarily repaired by the cells. For example, the upregulated FANCD2 (MCN FC: 4.3, MSN FC: 4.8) (Figure 6C) comes into the picture in response to DNA damage to participate in the repair process. FANCD2 forms complex with the RAD51 (MCN FC: 4.3, MSN FC: 1.5) during the S phase of the cell cycle during the DNA repair process by homologous recombination (Taniguchi et al., 2002).

The biological process term DNA replication (GO:0006260; MCN p-value 1.82E-04, MSN p-value 0.012962384) was found to be significantly downregulated that could potentially reflect DNA damage checkpoint (GO:0000077 MCN p-value 0.03872487, MSN p-value 0.046510499) activation observed in the GO analysis (Supplementary table 2) responsible for slowing down the DNA replicative machinery (Willis and Rhind, 2009). DNA checkpoints are activated in response to damage imposed on the cells to protect the cells from genotoxic stress and maintain a proper homeostasis (Verma et al., 2019).

TP53BP1, which functions as a DNA damage checkpoint protein, is shown to be upregulated (MCN FC: 5.6, MSN FC: 6.8) (Figure 6C) and is known to be involved in the DNA repair process by regulating the chromatin structure (DiTullio et al., 2002). It functions as a DNA damage checkpoint protein that functions in conjunction with ATM (MCN FC: 3.6459, MSN FC: 3.31609) (Mochan et al., 2004) that is also found to be enriched in our transcriptomics data and with subsequent enrichment of protein phosphorylation (GO:0006468; MCN p-value 0.002183143, MSN p-value 0.002006823). This might be suggestive of only initiation of the concerned pathways and not abrupt response. We can certainly provide this statement by considering the gene ontology data that shows more downregulated categories related to apoptosis than upregulated in the case of MCN and MSN (Supplementary table 2).

The cell cycle arrest in response to DNA damage (GO:0006977; MCN p-value 4.53E-04, MSN p-value 0.007577) was observed in the gene ontology category for both the NPs. The decreased DNA replication process also suggests the impact of DNA damage (GO:0006260: MCN p-value 1.82E-04, MSN p-value 0.013) and affected cell cycle process G2/M transition of the mitotic cell cycle (GO:0000086; MCN p-value 0.023, MSN p-value 0.0037) (Supplementary table 2). We found that the decrease in the G2/M transition process may signify the cellular machinery to prevent the abrupt responses due to NPs exposure. The decrease in the transition is also postulated by the decreased expression of various important regulators. Calmodulin (CALM) plays a significant role in metaphase transition or later phases of the cell cycle. It also stimulates the expression of various genes involved in the cell cycle progression or required by the cells for transition through G1 to mitosis (Rasmussen and Means, 1989). The decreased CALM (MCN FC: −11, MSN FC: −11.8) activity also resulted in G1 arrest and delays G2/M transition (Patel et al., 1999), which is also reflected in our RNA sequencing data by NPs response. We have also observed the downregulation of MELK (MCN FC: −13.1, MSN FC: −12.6), which is a cell cycle-dependent serine/threonine-protein kinase involved in cell cycle and DNA repair by recruiting repair proteins (Jiang and Zhang, 2013; Beke et al., 2015). The downregulation of the significant centrosomal gene; CEP70 (MCN FC: −6.8, MSN FC: −9.7) was observed that is required for the microtubule spindle assembly by binding to γ-tubulin (Shi et al., 2011). This might represent an interference with microtubule assembly by NPs that slows the progression of the mitosis phase of the cell cycle. It has also been seen that microtubule disassembly affects the G2/M transition phase, and cells delay division (Rieder and Cole, 2000).

#### 3.6 Modulated transcriptional machinery in response to mesoporous nanostructures

The exposure of NMs caused the stress that induced transcription repression by activating certain repressors to switch off specific genes. The most imp way to cope with the stress conditions inside the cells is to limit the expression of proteins. The cellular condition directly influences turnover rates of mRNA either by its decay or by modifying the interaction of various mRNP to achieve stability (Guzikowski et al., 2019). Transcriptional changes inside the cells in response to any external stressor can generate a cellular defence system. The enriched terms in the DAVID analysis, including negative regulation of transcription from RNA polymerase II promoter (GO:0000122) and negative regulation of transcription DNA templated (GO:0045892), suggest changing the milieu of the mRNA population inside the cell during NMs exposure. The cellular component categories nucleus (GO:0005634), nucleoplasm (GO:0005654) and transcription factor complex (GO:0005667) suggested perturbed transcriptional changes due to MSN exposure (Supplementary table 2). Venn diagram analysis revealed potential similarities among the NPs with 23% upregulated and 25.6% downregulated transcription related genes (Figure 7A).

**Figure 7:**
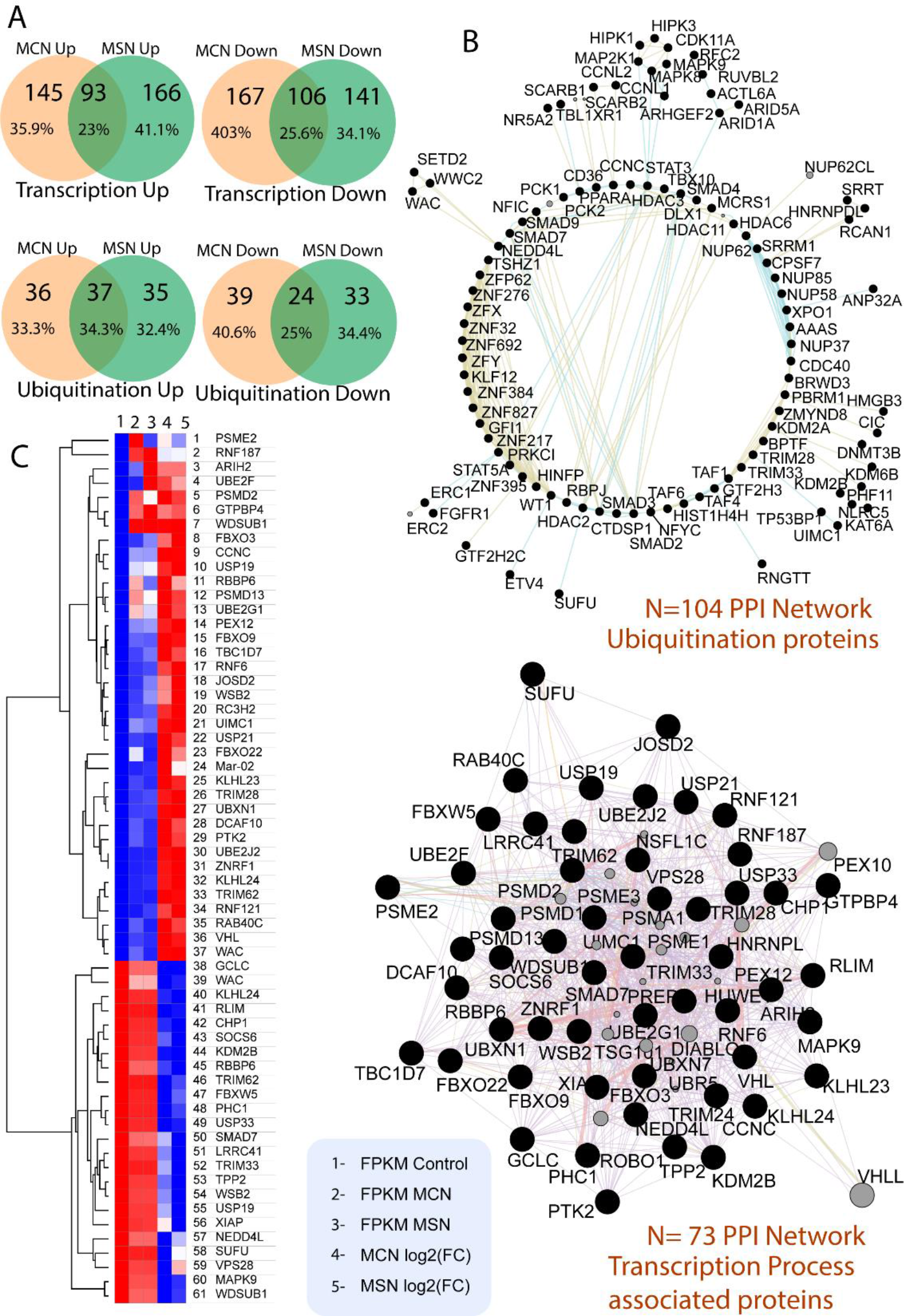
Transcription related gene alteration by mesoporous NPs: The Venn diagram represents unique and commonly shared deregulated genes of transcriptional machinery of the cell. (A) Represents upregulated genes regulated by MCN and MSN where 11.2% of the genes were common, 75.6% genes were regulated by MCN and 13.1% were regulated by MSN uniquely. MCN and MSN NPs commonly downregulated 22.3% genes whereas 37.9% genes by MCN and 39.8% genes by MSN NPs were uniquely downregulated. (C) Displaying heatmap of the genes detected in the overlap of the Venn diagrams regulated commonly by both the NPs. Columns are ordered by FPKM values of control, MCN and MSN NPs.

The NMs are also able to regulate deacetylases suggested by the GO categories histone deacetylation (GO:0016575) and protein deacetylation (GO:0006476) that plays a pivotal role in gene silencing as evidenced by the high upregulation of HDAC1 (MCN FC: 1.5, MSN FC: 10.4) (Supplementary figure 6). They might indicate that more transcriptional changes have been carried in the cells due to MSN exposure that do not necessarily activate stress-related genes but can resist more adverse perturbations.

The observed primary transcriptional response due to exposure to MSN and MCN can be a strategy of a cell to mitigate low levels of stress that ultimately limits the expression of other stressful pathways. We can explain this based on the results obtained from the cellular assay MTT, in which no cytotoxic effects were seen at the lower concentrations (15 μg/ml MSN) and initial periods (6 h), but more changes are occurring at the molecular levels even at non-cytotoxic doses. The high upregulation of specific genes can also suggest this scenario, including histone H4 transcription factor (HINFP; MCN FC: 10, MSN FC: 12), that is responsible for regulating the transcription of histone genes as well as other target genes such as those participating in DNA repair and cell cycle regulatory functions (Mitra et al., 2003). This can be perceived as a cellular defence strategy against the NPs insult. Furthermore, the enriched category related to apoptosis in MCN and MSN (Supplementary table 2) suggested that though NPs are trying to impose the toxic insult, the cellular system is trying to resist the harmful effects by limiting the aberrant uncontrolled proliferation of cells. This behaviour can also be explained by the highly upregulated expression of cell surface serine protease hepsin; HPN (MCN FC: 10 MSN FC: 11.8) responsible for maintaining proper cell morphology, proliferation state of the hepatoma cells (Torres et al., 1993).

The stability of mRNA largely depends upon the poly (A) tail that protects its integrity under different conditions. The role of cellular machinery in mediating the mRNA turnover rates during NMs exposure was suggested by the enrichment of GO term regulation of mRNA stability, nuclear-transcribed mRNA poly(A) tail shortening (GO:0000289) and subsequent high-level upregulation of Poly(A)-specific ribonuclease PARN (MCN FC 13.65, MSN FC 14.15). PARN mediates the 3’-exoribonuclease activity through efficient degradation of poly(A) tails of mRNA and prevents mRNA expression that may contain premature stop codons. The poly (A) dynamics changed during DNA damage, in which PARN played an important role (Cevher et al., 2010). This might result in silencing or degradation of aberrant mRNAs during stress conditions. Thus during the NMs exposure, mRNA decay levels may provide an additional backup strategy to deal with stress initiated conditions.

In contrast to MCN, more transcription related GO terms were found to be downregulated for MSN category (Supplementary table 2) that includes RNA processing (GO:0006396 p-value 5.40E-04), chromatin remodeling (GO:0006338 p-value 0.010131), mRNA export from nucleus (GO:0006404 p-value 0.013526), mRNA splicing, via spliceosome (GO:0000398 p-value 0.019923), regulation of mRNA stability (GO:0043488 p-value 0.037019), nuclear-transcribed mRNA poly(A) tail shortening (GO:0000289 p-value 0.038307), 3’-UTR-mediated mRNA destabilization (GO:0061158 p-value 0.039591), negative regulation of sequence-specific DNA binding transcription factor activity (GO:0043433, p-value 0.041205) and ~histone H3 deacetylation (GO:0070932, p-value 0.399361). Additionally, the common transcription related downregulated categories by both the NPs includes DNA replication, transcription DNA templated, regulation of transcription DNA templated and negative regulation of transcription from RNA polymerase II promoter (Supplementary table 2). It has been seen that during the exposure of stress by the cellular system, for instance, DNA damage, there is a possibility that NPs imposed RNA polymerase arrest to block the elongation that leads to initiation of DNA repair pathways that allows transcription-coupled repair (Yu et al., 2016). The transcription by the RNA polymerase II is tightly regulated by the signalling pathways (Chen et al., 2018). Importantly, during the stress and damage, cells slow down RNA Pol II’s progression (Wang et al., 2018). Number of transcription factors such as zinc finger nucleases (ZNF), SMAD genes (SMAD7 and SMAD9, SMAD3), HINFP, STAT3 have been perturbed in our RNA sequencing dataset interacting closely with each other to elicit a major response signified in the PPI network (Figure 7B).

#### 3.7 Ubiquitination mediated changes in the protein pool

Though transcriptional induction during stress conditions plays a pivotal role in the cellular defence system, posttranslational modifications depend upon altering protein structures through various forms, including ubiquitination, phosphorylation, methylation, acetylation, and sumoylation. This strategy allows a faster response against cellular stress by modifying pre-existing proteins that regulate various activities and pathways. The ubiquitination is a hallmark of posttranslational covalent modification that not only tags the proteins for degradation but also makes them work according to the cellular conditions. Our transcriptomic dataset found >200 upregulated and >100 downregulated transcripts involved in the ubiquitination pathway, indicating substantial protein modification during MCN and MSN exposure (Supplementary table 5). Furthermore, 34.3% of upregulated transcripts and 25% downregulated transcripts for ubiquitination were shared between MCN and MSN, which suggested almost similar biological response perturbation at an early low dose exposure (Figure 7A).

During the GO functional analysis, various biological processes and subsequent genes related to ubiquitination were enriched. For example, the GO category protein ubiquitination (GO:0016567; MCN p-value-3.78E-04, MSN p-value 0.003299) was significantly enriched for the MSN and MCN that mainly might regulate fundamental mechanisms during cellular stress. Also, the categories for genes consists of protein K48-linked ubiquitination (GO:0070936, MCN p-value 0.002472, MSN p-value 9.03E-04), ubiquitin-dependent protein catabolic process (GO:0006511, MCN p-value 0.005802, MSN p-value 0.01003) that suggested protein degradation due to NPs stress as ubiquitination on lysine 48 (K48) tagged the protein for degradation with proteasomal pathway.

We have also observed the upregulation of various genes of proteasomal subunits (Supplementary table 5) that also signifies cellular efforts against the NMs induced stress. This is also signified by the enriched BP category proteasome-mediated ubiquitin-dependent protein catabolic process (GO:0043161; MCN p-value 0.018735, MSN p-value 4.70E-04) and also from the PPI network (Figure 7D). The increased oxidative stress may partly attribute this by NPs increased the expression of stress response transcription factor Nrf1 or NFE2L1 (MCN FC: 1.5, MSN FC: 1.3), leading to enhanced transcription of proteasomal subunit genes in hepatocytes (Lee et al., 2013) and thus restoring the protein homeostasis.

Of crucial importance, the ubiquitination pathway orchestrates other cellular processes by controlling the turnover rates of proteins acting in that process to synchronise the harmony in the cell. For example, high upregulation of an E3 ubiquitin ligase RAB40C (CN 12.52, SN 10.94) was observed in high throughput data that controls the turnover rates of RACK1 protein downregulated by targeting them for proteasomal degradation.

According to our data, we previously described the high enrichment of GO categories related to the transcription in both the NPs. The cellular machinery dedicated itself to quickly respond to changing conditions inside it by timely regulating its gene expression. Thus, simultaneous action of ubiquitination with transcriptional cascade is the main factor for controlling the outcome of gene expression during NPs stress conditions. This can also be shown convincingly since we discovered significant downregulation of transcription-related activities and subsequent overexpression of ubiquitination-related processes (Supplementary table 2), which may offer insight into how ubiquitination regulates gene expression. For instance, the high downregulation of a crucial histone deacetylase (HDAC2; MCN FC: −11, MSN FC: −11) might result from stress created by NMs and subsequent ubiquitination followed by degradation through the proteasomal complex. The ubiquitination and subsequent proteasomal degradation of HDAC2 were observed in response to cigarette smoke extract (Kim et al., 2011). Therefore, this could be a part of regulating histone-modifying enzymes by the ubiquitin-proteasome system to precisely fine-tune the transcriptional activity against NMs. The von Hippel-Lindau (VHL) E3 ubiquitin ligase, which was found to be highly upregulated (MCN FC: 11.51, MSN FC: 11.01) in our study, annotated for the number of GO categories such as regulation of transcription (GO:0006355), positive regulation of transcription from RNA pol II promoters (GO:0045893), negative regulation of cell proliferation (GO:0008285) and protein ubiquitination that signifies its prominent role in regulating the number of processes.

In addition to upregulation, simultaneous downregulation of proteasomal ubiquitin-dependent protein catabolic process (GO:0032436) has also been observed for NMs response. The downregulation of predominant E3 ubiquitin ligase MDM2 (MCN FC: −2.4, MSN FC: −3.1) and upregulation of TP53 (MCN FC: 7, MSN FC: 1.8) are evidence of the critical interplay between ubiquitination pathways and cellular processes during external stress. Thus, the cells during NMs exposure might have regulated the low levels of ubiquitin ligase MDM2 to keep the TP53 levels high, resulting in implications on various pathways to maintain the homeostasis.

#### 3.8 Conclusion

Using the high-throughput technique, we have conducted a safety assessment of MCN and MSN for a future reference towards their biocompatibility assessment. The safety evaluation of mesoporous nanoparticles alone other than in combination with a drug helps predict any collateral damage to the biological system. Notably, we have observed that two different mesoporous nanoparticles triggered similar molecular responses at low exposure levels (Figure 8).

**Figure 8:**
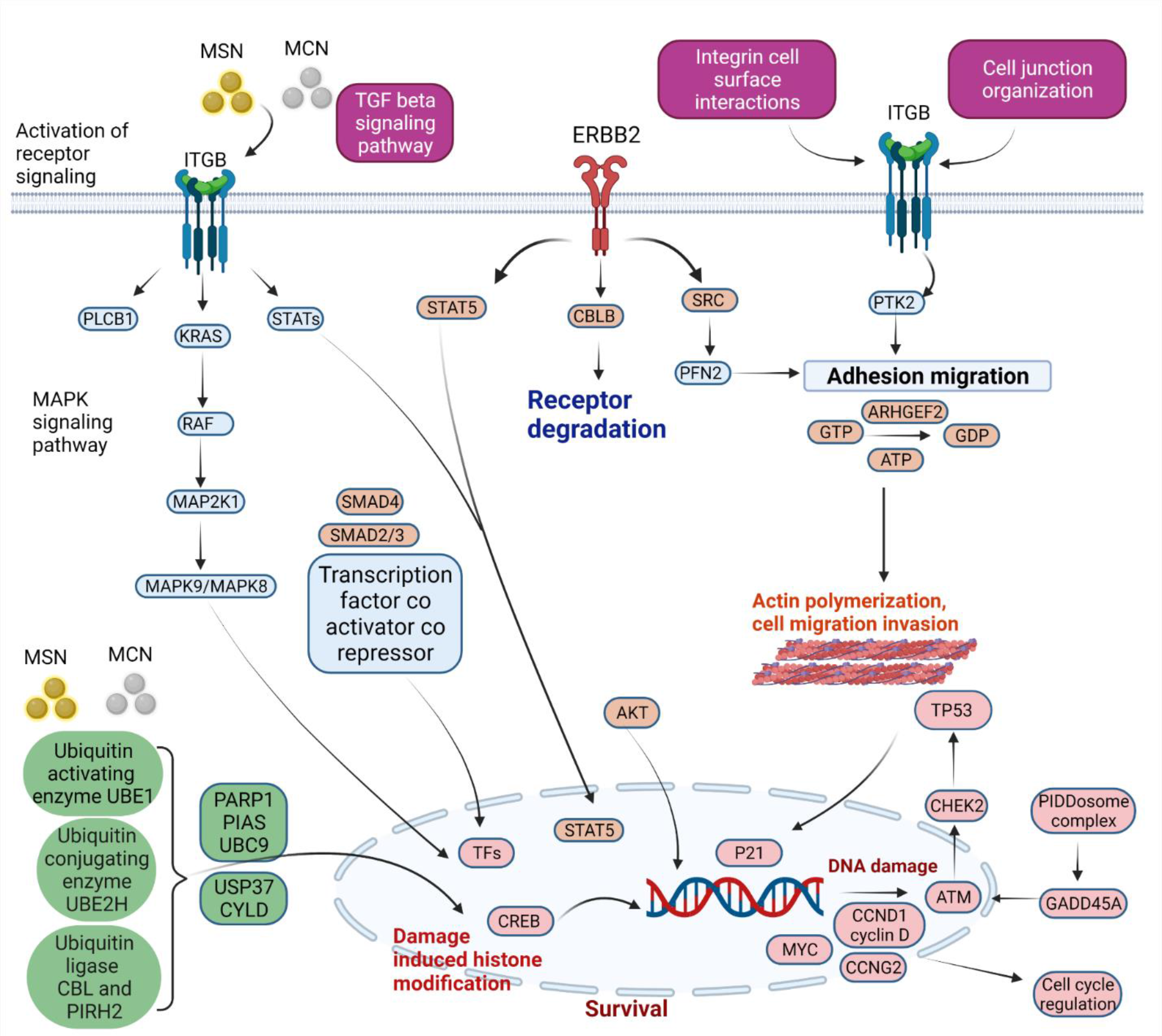
Schematic representation of the initiation of signaling pathways in response to mesoporous silica and mesoporous carbon nanoparticles

The cell viability data reveals dose and time-dependent effects of nanoparticles, but subsequently, a low dose was selected for further evaluation. The transcriptomics data identified initiated molecular responses that are not related to extremely harmful compared to the other toxicants. The gene expression analysis revealed that mesoporous nanostructures could exert molecular changes by the intensified changes in common gene ontology terms. With the in-depth gene ontology analysis, we observed that the cellular system responded with increased transcriptional activity by regulating mRNA decay, histone modifiers and transcription factor (HINFP) that regulated histone modification in addition to other regulatory mechanisms inside the cell. The dynamic interaction of MCN and MSN with cytoskeletal machinery affected actin and microtubule activity. The increased actin and microtubule activity indicated the internalisation of high aspect ratio nanoparticles. This is also supported by the information gathered during RNA sequencing data in which we identified processes related to the endocytosis, but we did not explain extensively.

Furthermore, increased expression of DNA damage and DNA checkpoint genes allowed initiation of DNA repair machinery that transiently stalls the replication process indicated by the downregulated G2/M transition and DNA replication ontology categories. We have also observed increased ubiquitination and proteasomal activity to alter the protein turnover to mitigate the stress condition imposed by mesoporous nanoparticles. The increased apoptosis-related ontology also sees the impact of DNA damage by the nanoparticles indicating cell death but not the harmful levels as it is balanced by the more enrichment of the downregulated apoptosis process. This could be due to the low exposure duration of the NPs that prevented the nano-bio interaction that could pose a more significant impact. The present study discloses the underlying molecular mechanism against the mesoporous nanoparticles, more importantly, increased understanding of nano-bio interaction, which should be of great significance during the efficient development of the drug delivery system.

